# Induction of AMPK activation by *N*,*N’*-Diarylurea FND-4b decreases growth and increases apoptosis in triple negative and estrogen-receptor positive breast cancers

**DOI:** 10.1101/489179

**Authors:** Jeremy Johnson, Piotr Rychahou, Vitaliy M. Sviripa, Heidi L. Weiss, Chunming Liu, David S. Watt, B. Mark Evers

## Abstract

**Purpose:** Triple negative breast cancer (TNBC) is the most lethal and aggressive subtype of breast cancer. AMP-activated protein kinase (AMPK) is a major energy regulator that suppresses tumor growth, and 1-(3-chloro-4-((trifluoromethyl)thio)phenyl)-3-(4-(trifluoromethoxy)phenyl)urea (FND-4b) is a novel AMPK activator that inhibits growth and induces apoptosis in colon cancer. The purpose of this project was to test the effects of FND-4b on AMPK activation, proliferation, and apoptosis in breast cancer with a particular emphasis on TNBC.

**Materials and methods:** (i) Estrogen-receptor positive breast cancer (ER+BC; MCF-7, and T-47D), TNBC (MDA-MB-231 and HCC-1806), and breast cancer stem cells were treated with FND-4b for 24h. Immunoblot analysis assessed AMPK, acetyl-CoA carboxylase (ACC), ribosomal protein S6, cyclin D1, and cleaved PARP. (ii) Sulforhodamine B growth assays were performed after treating ER+BC and TNBC cells with FND-4b for 72h. Proliferation was also assessed by counting cells after 72h of FND-4b treatment. (iii) Cell death ELISA assays were performed after treating ER+BC and TNBC cells with FND-4b for 72h.

**Results:** (i) FND-4b increased AMPK activation with concomitant decreases in ACC activity, phosphorylated S6, and cyclin D1 in all subtypes. (ii) FND-4b decreased proliferation in all cells, while dose-dependent growth decreases were found in ER+BC and TNBC. (iii) Increases in apoptosis were observed in ER+BC and the MDA-MB-231 cell line with FND-4b treatment.

**Conclusions:** Our findings indicate that FND-4b decreases proliferation for a variety of breast cancers by activating AMPK and has notable effects on TNBC. The growth reductions were mediated through decreases in fatty acid synthesis (ACC), mTOR signaling (S6), and cell cycle flux (cyclin D1). ER+BC cells were more susceptible to FND-4b-induced apoptosis, but MDA-MB-231 cells still underwent apoptosis with higher dose treatment. Further development of FND compounds could result in a novel therapeutic for TNBC.

## Introduction

Breast cancer is the most common cancer in women and the main cause of cancer-related death among women worldwide. In 2018 alone, there will be more than 266,000 newly diagnosed cases of breast cancer in women in the United States and almost 41,000 deaths [1]. Up to 30% of patients develop metastases, and 90% of deaths result from metastases to the lung, brain, or bone [2]. Breast cancer is a heterogeneous disease separable into three main types: estrogen-receptor positive breast cancer (ER+BC), HER2-amplified breast cancer, and triple negative breast cancer (TNBC). Although TNBC comprises only 15-20% of total cases, it is the most lethal and aggressive of the three types [3, 4].

The principal characteristics of TNBC include: (1) reduced expression of the estrogen and progesterone receptors and (2) no overexpression of HER2. TNBC affects a younger patient population than the population afflicted with other types of breast cancer and leads to an increased risk of recurrence and metastases [3]. Not surprisingly, patients with recurrent TNBC have a worse prognosis than that for patients with recurrent forms of other breast cancers [3]. In addition, patients with TNBC have limited therapeutic options because their tumors lack the traditional steroid hormone receptors and HER2 amplification. Instead, patients usually receive a drug cocktail that includes an anthracycline antineoplastic agent, a DNA alkylating agent, and a taxane [3]. These chemotherapeutic agents are toxic to normal and cancer cells alike and result in serious side-effects that are difficult for patients to tolerate. Recent efforts have focused on developing therapies that specifically target cancer cells without affecting normal cells. Because oncogenic transformation requires major metabolic reprogramming to produce energy, redox cofactors, and molecules involved in DNA modification, new agents that target the increased metabolism within cancer tissue more than the metabolism in normal tissue are attractive therapeutic options [2].

AMP-activated protein kinase (AMPK) is a cellular energy sensor that has important implications in cancer progression [5–16]. When activated by ATP depletion, the phosphorylated form of AMPK causes the following changes in TNBC: (1) inhibition of anabolic and oncogenic pathways, (2) attenuated mTOR signaling, (3) decreased cell proliferation, and (4) apoptosis [17–26]. Well-known AMPK activators, such as 5-aminoimidazole-4-carboxamide ribonucleotide (AICAR) and 2-deoxyglucose (2-DG), require high doses to affect cancer cell proliferation, which has led to their unsuccessful translation to the clinic for cancer therapy [12]. Among the attempts to produce new AMPK activators with increased sensitivity, the fluorinated *N*,*N’*-diarylureas (FNDs) serve as activators that lead to phosphorylated AMPK at low concentrations [27, 28]. In particular, 1-(3-chloro-4-((trifluoromethyl)thio)phenyl)-3-(4-(trifluoromethoxy)phenyl)urea (FND-4b) inhibits growth and induces apoptosis in colorectal cancer cells, but its potential effects on other types of cancer remain unclear [27]. Because of the pressing need to develop new treatments for TNBC, we tested the effects of FND-4b on TNBC and compared the results with ER+BC. Importantly, we found that treatment with FND-4b led to AMPK activation, decreased cell cycle flux, and increased apoptosis in both subtypes. These findings indicate that FND compounds may be potential therapeutic options for TNBC.

## Materials and methods

### Reagents, supplements, and antibodies

1-(3-Chloro-4-((trifluoromethyl)thio)phenyl)-3-(4-(trifluoromethoxy)phenyl)urea (FND-4b) was synthesized and characterized as previously described [28]. Roswell Park Memorial Institute (RPMI) 1640 Medium and Eagle’s Minimum Essential Medium (EMEM) were purchased from Sigma-Aldrich (St. Louis, MO). Dulbecco’s Modified Eagle Medium (DMEM) was from Corning (Corning, NY). Human breast cancer stem cell complete growth medium was from Celprogen (Torrance, CA). MEM non-essential amino acid solution (100x), sodium pyruvate solution (100 mM), insulin solution (10 mg/mL), penicillin-streptomycin (100x) (P/S), and fetal bovine serum (FBS) were from Sigma-Aldrich. Antibodies for pAMPKα (Thr172), total AMPKα, phosphorylated acetyl-CoA carboxylase (ACC), total ACC, phosphorylated ribosomal protein S6, total S6, and PARP were from Cell Signaling Technology (Danvers, MA). The cyclin D1 antibody was from Abcam (Cambridge, MA). The beta-actin antibody was from Sigma-Aldrich. The Sulforhodamine B (SRB) Cytotoxicity Assay was from G-Biosciences (St. Louis, MO). The Cell Death Detection ELISA^PLUS^ assay was from Sigma-Aldrich.

### Cell culture

MCF-7, T-47D, MDA-MB-231, HCC-1143, and HCC-1806 cells were purchased from ATCC, while breast cancer stem cells were purchased from Celprogen. MCF-7 cells were maintained in EMEM containing 10% FBS, 1% P/S, 0.01 mg/mL insulin, 1x non-essential amino acids, and 1 mM sodium pyruvate. T-47D cells were maintained in RPMI containing 10% FBS, 1% P/S, and 0.2 Units/mL insulin. MDA-MB-231, HCC-1143, and HCC-1806 cells were maintained in RPMI with 10% FBS and 1% P/S. Breast cancer stem cells were maintained in breast cancer stem cell medium supplemented with 10% FBS and 1% P/S. All cells were grown in an incubator at 37°C and 5% CO_2_. For cell treatments, 7×10^5^ MCF-7 and T-47D cells or 8×10^5^ MDA-MB-231, HCC-1806, and breast cancer stem cells were seeded in 6-well plates and incubated overnight. The medium was removed on the following day, and cells were treated with fresh medium that contained different concentrations of FND-4b (0, 1, 2.5, 5, 10, and 20 μM) for 24 h before lysis.

### Western blot analysis

After treatment, cells were scraped from the wells with 1x RIPA buffer containing serine protease inhibitor. The cells were lysed by incubating on ice for 20 min and vortexing 10 sec every 5 min. The lysates were centrifuged at 14,000 rpm and 4°C for 20 min, and the protein concentration was determined using the Bradford method. The proteins were reduced and denatured by heating at 80°C for 10 min. An equal amount of protein was resolved on SDS-PAGE gels and transferred to PVDF membranes. The membranes were blocked with 10% milk before overnight incubation with primary antibodies at 4°C. On the following day, the membranes were washed with Tris buffered saline with 0.1% tween-20 (TBST) for 5 min and again for 10 min. The membranes were subsequently incubated with the appropriate secondary antibody for 30 min at room temperature. The membranes were then washed with TBST for 15 min and again for 20 min. Proteins were visualized with Amersham ECL (GE Healthcare) or Immobilon (Millipore). Membranes were stripped and reprobed as necessary.

### Cell counting assay

Cells (1×10^5^) were seeded in 6-well plates and incubated overnight. The medium was removed on the following day, and cells were grown in fresh medium that contained 0 or 5 μM FND-4b for 72 h. Cells were then washed with PBS, trypsinized, and counted with a Beckman-Coulter cell counter.

### SRB growth assay

Cells (5×10^3^ in 100 μL) were seeded in 96-well plates and incubated overnight. On the following day, fresh media was prepared to contain twice the desired concentrations of FND-4b (i.e., 5, 10, 20, and 40 μM). Then 100 μL of the new media solutions were added to the wells without removing the old media; this yielded 200 μL per well and halved the FND-4b concentrations. Cells were grown in the media with different final concentrations of FND-4b (0, 2.5, 5, 10, and 20 μM) for 72 h before the proteins were fixed at 4°C for 1 h. The wells were washed 3 times with water and then dried for 30 min. The SRB dye solution (0.4%) was added to the wells and incubated for 30 min. The excess dye was washed off with 1% acetic acid, and the wells were allowed to air dry. The dye was solubilized with a 10 mM Tris solution, and the absorbance was measured at 565 nm or—if the readings were outside of the instrument’s linear range—at 490 nm.

### Cell Death Detection ELISA^PLUS^ assay

Cells (5×10^3^ in 100 μL) were seeded in 96-well plates and incubated overnight. On the following day, fresh media was prepared to contain twice the desired concentrations of FND-4b (i.e., 5, 10, and 20 μM). Then 100 μL of the new media solutions were added to the wells without removing the old media; this yielded 200 μL per well and halved the FND-4b concentrations. Cells were cultured in the media with different final concentrations of FND-4b (0, 2.5, 5, and 10 μM) for 72 h. Cells were then centrifuged at 200 x g for 10 min and lysed for 30 min with shaking. The lysates were centrifuged at 200 x g for 10 min, and 20 μL of supernatant was transferred to streptavidin-coated wells. Then 80 μL of the immunoreagent (anti-histone biotin and anti-DNA peroxidase) was added to the streptavidin-coated wells. The plates were shaken for 2 h at room temperature before the wells were rinsed three times with incubation buffer. Color change was initiated by adding the substrate ABTS to the wells, and the plates were shaken until the color change was sufficient for photometric analysis. After adding the ABTS Stop Solution, absorbance was measured at 405 nm.

### Statistical analysis

Comparisons of SRB growth and ELISA assays across non-treated and different dose groups of FND-4b were performed using analysis of variance (ANOVA) with test for linear trend across dose levels. Pairwise comparisons of each FND-4b dose level versus non-treated group were performed within the ANOVA model with Holm’s p-value adjustment for multiple testing. Comparisons of cell counting assays between control and FND-4b-treated groups were performed using two-sample t-tests with homogeneity of variance assessed for the use of the t-test for two group comparisons. Analyses were performed on data normalized with the non-treated group for the SRB growth and cell counting assays and on the raw data for the ELISA assays. In all experiments, p-values less than 0.05 were considered significant.

## Results

### Analysis of pAMPKα expression in breast cancer subtypes

Prior work has suggested that TNBC cell lines and tissues have higher levels of phosphorylated and total forms of AMPKα than non-TNBC cells and tissues [29]. Consequently, we compared levels of pAMPKα and total AMPKα in TNBC and ER+BC cells using western blotting. We found no difference in levels of phosphorylated or total AMPKα between two ER+BC cell lines (MCF-7 and T-47D) and three TNBC cell lines (MDA-MB-231, HCC-1143, and HCC-1806) (**Fig 1**).

**Fig 1.**
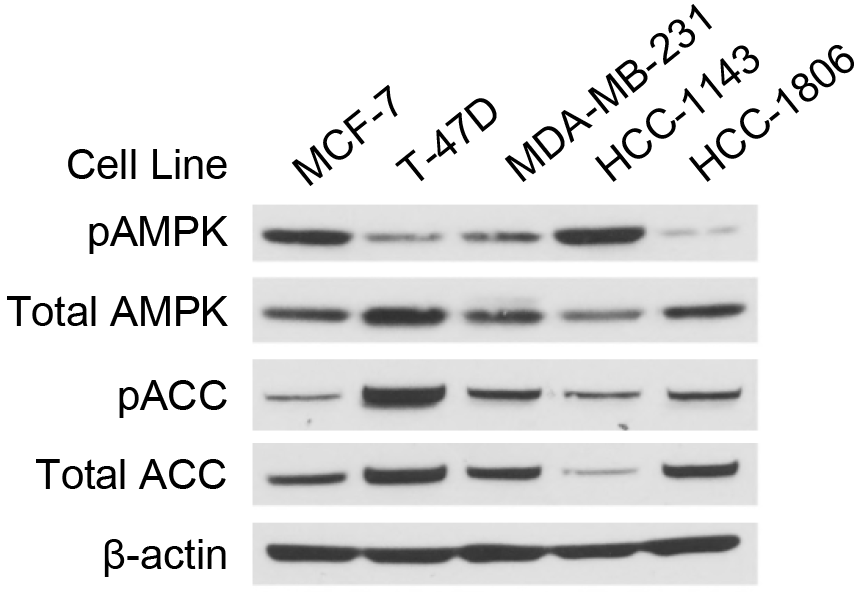
Examination of pAMPKα levels in breast cancer subtypes. Equal numbers of MCF-7, T-47D, HCC-1143, MDA-MB-231, and HCC-1806 cells were incubated overnight before cell lysis. Western blot analyses were performed for total and phosphorylated forms of AMPKα and ACC. Beta-actin was used as the loading control.

### FND-4b activated AMPKα and downstream signaling pathways in breast cancer

Because FND-4b activated AMPKα in colon cancer cells, we investigated its effects in several subtypes of breast cancer [27]. In particular, MCF-7, T-47D, MDA-MB-231, HCC-1806, and breast cancer stem cells were treated with fresh medium containing a range of FND-4b concentrations (0, 1, 2.5, 5, 10, and 20 μM) for 24 h. Protein levels of phosphorylated and total forms of AMPKα, acetyl-CoA carboxylase (ACC), and ribosomal protein S6, cyclin D1, and cleaved PARP were measured with immunoblotting. ACC and S6 were selected for analysis because they are downstream effectors of AMPKα that are phosphorylated and dephosphorylated, respectively, with AMPKα activation. As expected based on prior work, FND-4b treatment increased levels of pAMPKα in all five cell lines (**Fig 2**) [27]. Consistent with this result, increases in pACC and decreases in pS6 were also noted. Cyclin D1, an AMPKα-regulated marker for flux through the cell cycle, was decreased in all cell lines treated with FND-4b. Increases in the apoptotic indicator cleaved PARP were observed in MCF-7, MDA-MB-231, and HCC-1806 cells with AMPKα activation. FND-4b concentrations less than 5 μM yielded minimal effects on AMPK signaling, but the 5 μM dose yielded robust effects in all cells tested. The HCC-1806 cells were notable because AMPKα activation spiked with 5 μM FND-4b treatment and then declined with concentrations higher than 5 μM. Taken together, these results indicate that FND-4b activates AMPKα and its downstream signaling pathways in TNBC, ER+BC, and breast cancer stem cells.

**Fig 2.**
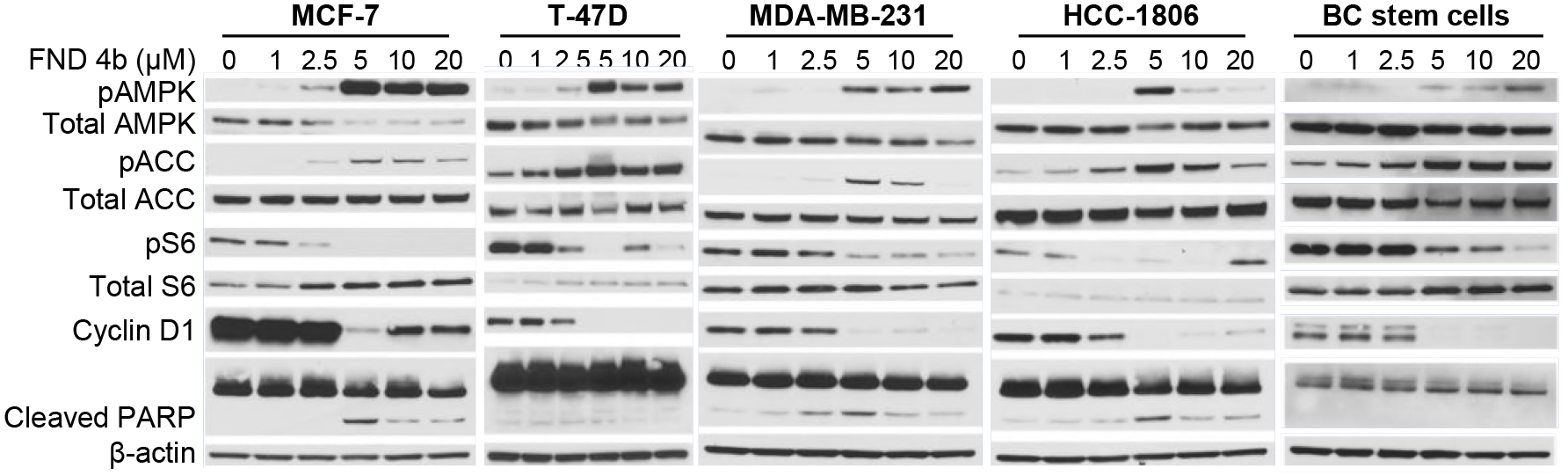
FND-4b activated AMPKα and its downstream signaling pathways in a dose-dependent manner in breast cancer. Equal numbers of MCF-7, T-47D, MDA-MB-231, HCC-1806 or breast cancer stem cells were treated with fresh medium that contained different FND-4b concentrations (0, 1, 2.5, 5, 10, and 20 μM) for 24 h. Western blot analysis was then performed for phosphorylated and total forms of AMPKα, ACC, and S6 as well as cyclin D1 and cleaved PARP. Beta-actin was used as the loading control. Treatment of the HCC-1806 cell line was repeated to confirm the spike in AMPKα activation at the 5 μM dosage with subsequent decreases at higher concentrations.

### FND-4b decreased growth of breast cancer subtypes in a dose-dependent fashion

AMPKα has been implicated as a tumor suppressor in breast cancer [17–26]. Since FND-4b activated AMPKα, we measured its effects on the growth of breast cancer cells. Cell counting assays showed that treatment at 5 μM FND-4b for 72 h resulted in significant growth inhibition of MCF-7, T-47D, MDA-MB-231, HCC-1806, and breast cancer stem cells (**Fig 3A**). Similar decreases occurred in all breast cancer subtypes with 5 μM treatment. In addition, SRB growth assays indicated that treatment at various FND-4b concentrations (2.5, 5, 10, and 20 μM) for 72 h yielded significant dose-dependent decreases in proliferation of MCF-7, T-47D, MDA-MB-231, and HCC-1806 cells (**Fig 3B**). ER+BC cells were more sensitive than TNBC cells to FND-4b at 2.5 μM, but the reductions were similar at higher concentrations. Similar growth inhibition at the 5 μM dosage between ER+BC and TNBC is consistent with the results from the cell counting assays. Taken together, these results illustrate that activation of AMPKα with FND-4b resulted in dose-dependent decreases in growth in ER+BC and TNBC.

**Fig 3.**
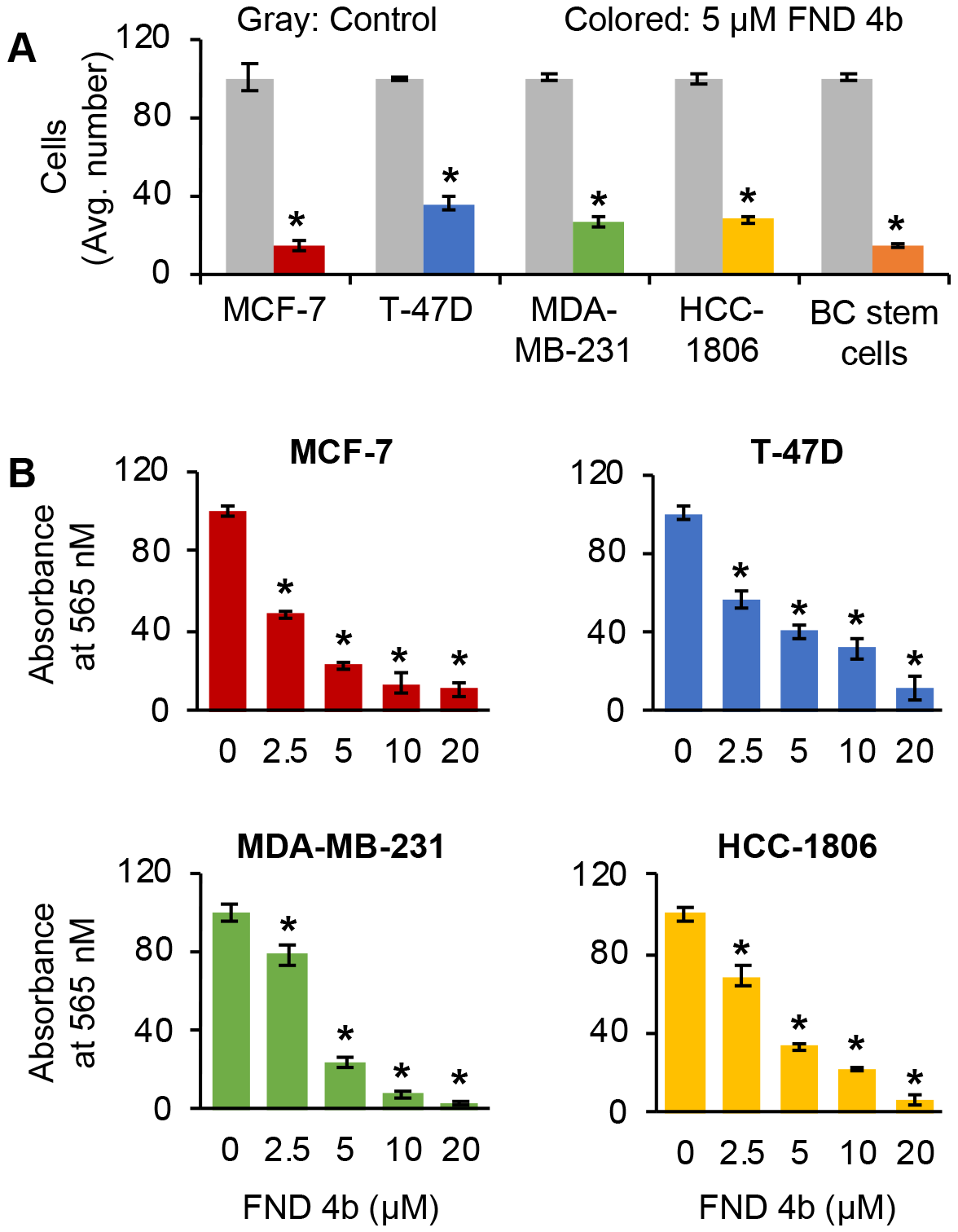
AMPKα activation with FND-4b decreased proliferation of breast cancer cells. (A) Equal numbers of MCF-7, T-47D, MDA-MB-231, HCC-1806, or breast cancer stem cells were grown in medium containing 0 or 5 μM FND-4b for 72 h followed by cell counting. (B) MCF-7, T-47D, MDA-MB-231, and HCC-1806 cells were grown in medium containing different concentrations of FND-4b (0, 2.5, 5, 10, and 20 μM) for 72 h before SRB growth assays were performed. Data are presented as mean ± SD from experiments performed in triplicate (cell counting) or sextuplicate (SRB assays) and are representative of three independent experiments. * indicates p-value < 0.001.

### FND-4b increased apoptosis in ER+BC and TNBC

While FND-4b’s major effect is on cell growth, its treatment also increased apoptosis in colon cancer [27]. Therefore, we investigated apoptosis induction in breast cancer cells. As previously mentioned, treatment with FND-4b resulted in increased levels of cleaved PARP, a marker of apoptosis, in MDA-MB-231, HCC-1806, and MCF-7 cells (see **Fig 2**). We also measured apoptosis with ELISA cell death assays that are more sensitive than western blotting assays. Significant increases in apoptosis were found in MCF-7 and T-47D cells—with MCF-7 cells being more sensitive (**Fig 4A-B**). Apoptosis was significantly increased in MDA-MB-231 cells with treatment at 10 μM FND-4b, but apoptosis in HCC-1806 cells was not increased (**Fig 4C-D**).

**Fig 4.**
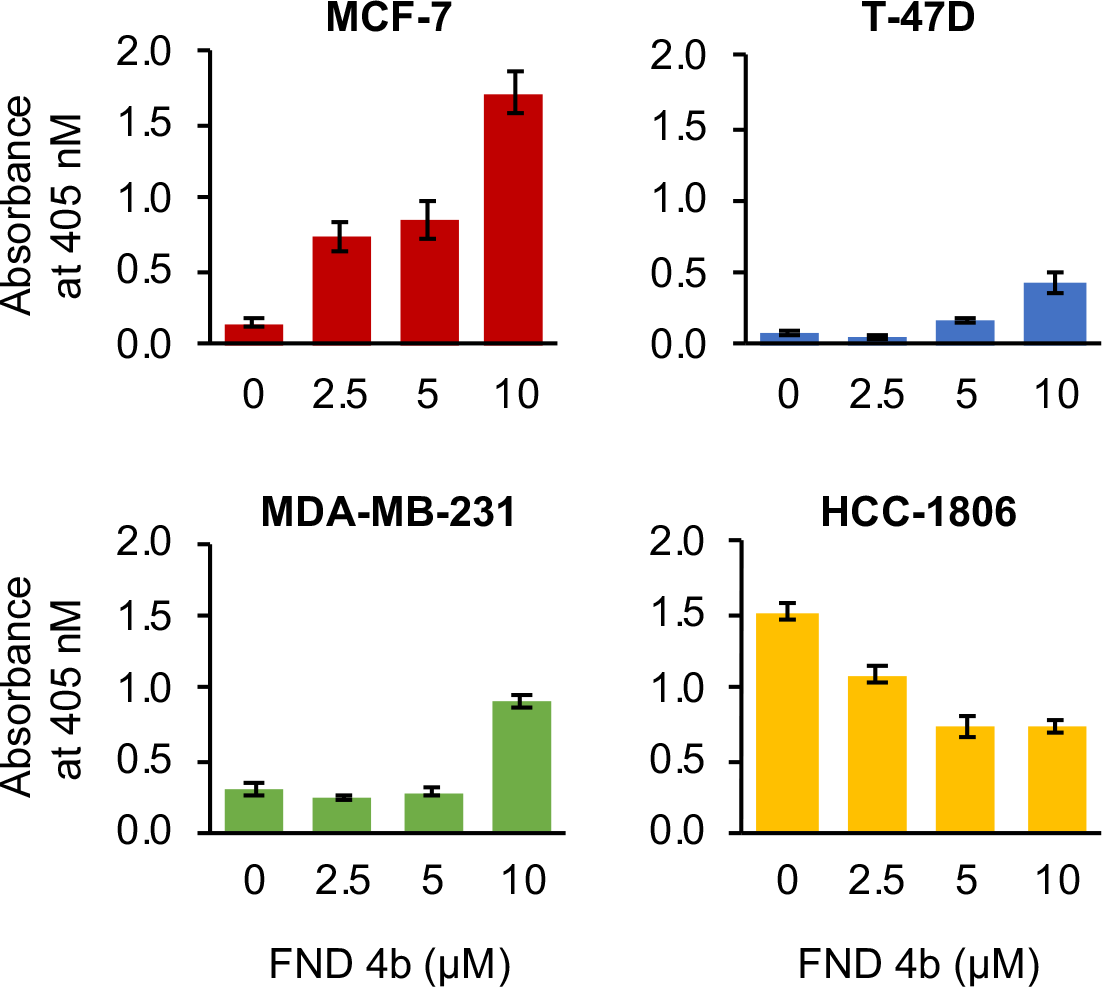
AMPKα activation with FND-4b increased apoptosis of breast cancer cells. Equal numbers of MCF-7, T-47D, MDA-MB-231, or HCC-1806 cells were cultured in medium containing different concentrations of FND-4b (0, 2.5, 5, and 10 μM) for 72 h before ELISA cell death assays were performed. Data are presented as mean ± SD from an experiment performed in triplicate and are representative of three independent experiments. * indicates p-value < 0.001.

## Discussion

The connection between AMPK activation and the inhibition of cancer cell growth prompted our interest in targeting AMPK and its downstream signaling pathways. AMPK activators such as AICAR and 2-DG have limited utility for patient care due to their high dose requirements [12]. We focused on novel small-molecule agents that activated AMPK at low concentrations and that stood a greater chance than these well-known AMPK activators of progressing toward the clinic. In this project, we examined the effects of the AMPK activator FND-4b to determine its effects on ER+BC, TNBC, and breast cancer stem cells. FND-4b previously suppressed the growth of colorectal cancer cells and stem cells through activation of AMPK in the low micromolar range without affecting signaling through the Akt or ERK pathways [27, 28]. We found that treatment with FND-4b resulted in dose-dependent increases in AMPK activation in both breast cancer subtypes and in the stem cells.

Other investigators have also focused on discovering novel AMPK activators or repurposing current drugs that activate AMPK for breast cancer therapy. OSU-53 and RL-71 activated AMPK and exerted anti-tumor effects in TNBC at low micromolar doses [17, 19]. In addition, demethoxycurcumin (20 μM) resulted in AMPK activation and large decreases in TNBC and ER+BC cell proliferation [24]. Finally, treatment of TNBC with the anti-depressant fluoxetine (0.5 μM) caused AMPK activation and substantial reductions in cellular viability [25]. Taken together, these studies and ours suggest the merit in targeting the AMPK signaling pathway for the treatment of breast cancer.

We found that low micromolar concentrations of FND-4b substantially reduced the growth of TNBC. This is particularly important because of the inherent aggressiveness of TNBC that has higher rates of recurrence and metastasis than other breast cancer subtypes [3]. The seriousness of TNBC is further amplified by the fact that most breast cancer deaths result from metastatic lesions [2]. The difficulty in developing treatments for TNBC is due to the lack of the estrogen receptor and HER2 amplification. Drugs that target these proteins, such as trastuzumab and tamoxifen, are ineffective in TNBC. Instead, patients with TNBC typically receive a drug cocktail that damages normal cells in addition to the tumor and leads to significant side effects. Current efforts are focused on developing drugs that specifically target proteins or pathways that are exclusively altered in TNBC. Since expression of AMPK and pAMPK is lower in breast cancer than normal tissue, this signaling pathway attracted our attention as an option for targeted therapy [30–32].

AMPK activation resulted in substantial decreases in cell proliferation in all breast cancer subtypes that were tested. These reductions in growth were due to AMPK’s ability to regulate the cell cycle both directly and indirectly. Directly, AMPK activation can attenuate levels of cyclin D1, an important protein that controls cell cycle arrest during the G1 phase [33]. Prior work in ovarian cancer has suggested that AMPK activation causes degradation of cyclin D1 through a pathway involving glycogen synthase kinase 3β [33]. Once cyclin D1 is degraded, cells are prevented from progressing past the G1 phase [33]. We found that FND-4b-induced AMPK activation resulted in substantial decreases in cyclin D1 expression, resulting in cell cycle arrest. Indirectly, AMPK activation can affect the cell cycle through effects on mTOR and cellular metabolism. AMPK downregulates flux through the mTOR pathway, which can control the cell cycle through its downstream effectors S6 kinase 1 (S6K1) and eukaryotic translation initiation factor 4E-binding protein 1 [34]. In our study, we measured mTOR activity by blotting for levels of phosphorylated ribosomal protein S6, which is downstream from S6K1. We found decreases in S6 phosphorylation with FND-4b-induced AMPK activation, indicating less mTOR flux and cell cycle progression. Additionally, AMPK affects cell metabolism by phosphorylating and inhibiting ACC, which is the rate-limiting step in fatty acid synthesis. As a result, *de novo* lipogenesis is inhibited. Fatty acids are required for progression through the cell cycle—notably, during the G_1_-S and G_2_-M phases—and in their absence, cells will be unable to complete mitosis [16]. Instead they will be arrested at the G_2_-M checkpoint [16]. We showed that FND-4b-induced AMPK activation led to increased ACC phosphorylation, signifying less fatty acid synthesis and flux through the cell cycle.

In addition to inducing cell cycle arrest, AMPK can also act as a tumor suppressor by causing apoptosis [32]. We found that FND-4b causes apoptosis in a dose-dependent manner in ER+BC cells with ELISA cell death assays. We also showed clear increases in levels of cleaved PARP—an apoptotic indicator—in MCF-7 cells. TNBC cells were more resistant to apoptosis from FND-4b, but there was apoptosis in MDA-MB-231 cells with 10 μM treatment. Additionally, we found increases in cleaved PARP in MDA-MB-231 and HCC-1806 cells with western blot. Taken together, these data indicate that ER+BC cells are more susceptible to FND-4b-induced apoptosis than TNBC. However, as suggested in previous work, the effects of FND-4b on cell cycle progression are more pronounced and consistent than on apoptosis [27].

## Conclusions

We have shown that the novel compound FND-4b can activate AMPK in ER+BC, TNBC, and breast cancer stem cells. In addition, treatment with this compound can dose-dependently decrease proliferation and increase apoptosis in breast cancer cells. The effects on cellular growth are mediated via decreased cell cycle flux—as evidenced by reductions in cyclin D1 levels—and suppression of fatty acid synthesis and mTOR signaling. With such profound effects on proliferation, further development of FND compounds could lead to their inclusion in TNBC treatment regimens.

## Acknowledgements

The Markey Cancer Center’s Research Communications Office assisted with manuscript preparation, and the Biostatistics and Bioinformatics Shared Resource Facility of the University of Kentucky Markey Cancer Center performed statistical analysis.

## SUPPORTING INFORMATION

Data for SRB growth assays, cell proliferation assays and ELISA assays. Primary antibody information.

## References

1. American Cancer Society. Cancer Facts & Figures 2018. Atlanta: American Cancer Society; 2018.

2. Elia I, Schmieder R, Christen S, Fendt SM. Organ-Specific Cancer Metabolism and Its Potential for Therapy. Handbook of experimental pharmacology. 2016;233:321–53. Epub 2015/04/29. doi: 10.1007/164_2015_10. PubMed PMID: 25912014.

3. Andreopoulou E, Schweber SJ, Sparano JA, McDaid HM. Therapies for triple negative breast cancer. Expert Opin Pharmacother. 2015;16(7):983–98. doi: 10.1517/14656566.2015.1032246. PubMed PMID: 25881743; PubMed Central PMCID: PMC5995333.

4. Yao H, He G, Yan S, Chen C, Song L, Rosol TJ, et al. Triple-negative breast cancer: is there a treatment on the horizon? Oncotarget. 2017;8(1):1913–24. doi: 10.18632/oncotarget.12284. PubMed PMID: 27765921; PubMed Central PMCID: PMC5352107.

5. Faubert B, Vincent EE, Poffenberger MC, Jones RG. The AMP-activated protein kinase (AMPK) and cancer: many faces of a metabolic regulator. Cancer Lett. 2015;356(2 Pt A):165–70. doi: 10.1016/j.canlet.2014.01.018. PubMed PMID: 24486219.

6. Grahame Hardie D. AMP-activated protein kinase: a key regulator of energy balance with many roles in human disease. J Intern Med. 2014;276(6):543–59. doi: 10.1111/joim.12268. PubMed PMID: 24824502; PubMed Central PMCID: PMC5705060.

7. Hardie DG, Alessi DR. LKB1 and AMPK and the cancer-metabolism link - ten years after. BMC Biol. 2013;11:36. doi: 10.1186/1741-7007-11-36. PubMed PMID: 23587167; PubMed Central PMCID: PMC3626889.

8. Hardie DG, Ross FA, Hawley SA. AMP-activated protein kinase: a target for drugs both ancient and modern. Chem Biol. 2012;19(10):1222–36. doi: 10.1016/j.chembiol.2012.08.019. PubMed PMID: 23102217; PubMed Central PMCID: PMC5722193.

9. Jeon SM. Regulation and function of AMPK in physiology and diseases. Exp Mol Med. 2016;48(7):e245. doi: 10.1038/emm.2016.81. PubMed PMID: 27416781; PubMed Central PMCID: PMC4973318.

10. Lage R, Dieguez C, Vidal-Puig A, Lopez M. AMPK: a metabolic gauge regulating whole-body energy homeostasis. Trends Mol Med. 2008;14(12):539–49. doi: 10.1016/j.molmed.2008.09.007. PubMed PMID: 18977694.

11. Li W, Saud SM, Young MR, Chen G, Hua B. Targeting AMPK for cancer prevention and treatment. Oncotarget. 2015;6(10):7365–78. doi: 10.18632/oncotarget.3629. PubMed PMID: 25812084; PubMed Central PMCID: PMC4480686.

12. Luo Z, Zang M, Guo W. AMPK as a metabolic tumor suppressor: control of metabolism and cell growth. Future Oncol. 2010;6(3):457–70. doi: 10.2217/fon.09.174. PubMed PMID: 20222801; PubMed Central PMCID: PMC2854547.

13. Motoshima H, Goldstein BJ, Igata M, Araki E. AMPK and cell proliferation--AMPK as a therapeutic target for atherosclerosis and cancer. J Physiol. 2006;574(Pt 1):63–71. doi: 10.1113/jphysiol.2006.108324. PubMed PMID: 16613876; PubMed Central PMCID: PMC1817805.

14. O’Neill HM, Holloway GP, Steinberg GR. AMPK regulation of fatty acid metabolism and mitochondrial biogenesis: implications for obesity. Mol Cell Endocrinol. 2013;366(2):135–51. doi: 10.1016/j.mce.2012.06.019. PubMed PMID: 22750049.

15. Sanli T, Steinberg GR, Singh G, Tsakiridis T. AMP-activated protein kinase (AMPK) beyond metabolism: a novel genomic stress sensor participating in the DNA damage response pathway. Cancer Biol Ther. 2014;15(2):156–69. doi: 10.4161/cbt.26726. PubMed PMID: 24100703; PubMed Central PMCID: PMC3928130.

16. Zadra G, Batista JL, Loda M. Dissecting the Dual Role of AMPK in Cancer: From Experimental to Human Studies. Mol Cancer Res. 2015;13(7):1059–72. doi: 10.1158/1541-7786.MCR-15-0068. PubMed PMID: 25956158; PubMed Central PMCID: PMC4504770.

17. Gao J, Fan M, Peng S, Zhang M, Xiang G, Li X, et al. Small-molecule RL71-triggered excessive autophagic cell death as a potential therapeutic strategy in triple-negative breast cancer. Cell Death Dis. 2017;8(9):e3049. doi: 10.1038/cddis.2017.444. PubMed PMID: 28906486; PubMed Central PMCID: PMC5636988.

18. Kim DE, Kim Y, Cho DH, Jeong SY, Kim SB, Suh N, et al. Raloxifene induces autophagy-dependent cell death in breast cancer cells via the activation of AMP-activated protein kinase. Mol Cells. 2015;38(2):138–44. doi: 10.14348/molcells.2015.2193. PubMed PMID: 25537862; PubMed Central PMCID: PMC4332026.

19. Lee KH, Hsu EC, Guh JH, Yang HC, Wang D, Kulp SK, et al. Targeting energy metabolic and oncogenic signaling pathways in triple-negative breast cancer by a novel adenosine monophosphate-activated protein kinase (AMPK) activator. J Biol Chem. 2011;286(45):39247–58. doi: 10.1074/jbc.M111.264598. PubMed PMID: 21917926; PubMed Central PMCID: PMC3234749.

20. Liu B, Fan Z, Edgerton SM, Deng XS, Alimova IN, Lind SE, et al. Metformin induces unique biological and molecular responses in triple negative breast cancer cells. Cell Cycle. 2009;8(13):2031–40. doi: 10.4161/cc.8.13.8814. PubMed PMID: 19440038.

21. Daurio NA, Tuttle SW, Worth AJ, Song EY, Davis JM, Snyder NW, et al. AMPK Activation and Metabolic Reprogramming by Tamoxifen through Estrogen Receptor-Independent Mechanisms Suggests New Uses for This Therapeutic Modality in Cancer Treatment. Cancer Res. 2016;76(11):3295–306. doi: 10.1158/0008-5472.CAN-15-2197. PubMed PMID: 27020861; PubMed Central PMCID: PMC4895922.

22. Gollavilli PN, Kanugula AK, Koyyada R, Karnewar S, Neeli PK, Kotamraju S. AMPK inhibits MTDH expression via GSK3beta and SIRT1 activation: potential role in triple negative breast cancer cell proliferation. FEBS J. 2015;282(20):3971–85. doi: 10.1111/febs.13391. PubMed PMID: 26236947.

23. Liu H, Scholz C, Zang C, Schefe JH, Habbel P, Regierer AC, et al. Metformin and the mTOR inhibitor everolimus (RAD001) sensitize breast cancer cells to the cytotoxic effect of chemotherapeutic drugs in vitro. Anticancer Res. 2012;32(5):1627–37. PubMed PMID: 22593441.

24. Shieh JM, Chen YC, Lin YC, Lin JN, Chen WC, Chen YY, et al. Demethoxycurcumin inhibits energy metabolic and oncogenic signaling pathways through AMPK activation in triple-negative breast cancer cells. J Agric Food Chem. 2013;61(26):6366–75. doi: 10.1021/jf4012455. PubMed PMID: 23777448.

25. Sun D, Zhu L, Zhao Y, Jiang Y, Chen L, Yu Y, et al. Fluoxetine induces autophagic cell death via eEF2K-AMPK-mTOR-ULK complex axis in triple negative breast cancer. Cell Prolif. 2018;51(2):e12402. doi: 10.1111/cpr.12402. PubMed PMID: 29094413.

26. Hadad SM, Hardie DG, Appleyard V, Thompson AM. Effects of metformin on breast cancer cell proliferation, the AMPK pathway and the cell cycle. Clin Transl Oncol. 2014;16(8):746–52. doi: 10.1007/s12094-013-1144-8. PubMed PMID: 24338509.

27. Kenlan DE, Rychahou P, Sviripa VM, Weiss HL, Liu C, Watt DS, et al. Fluorinated N,N’-Diarylureas As Novel Therapeutic Agents Against Cancer Stem Cells. Mol Cancer Ther. 2017;16(5):831–7. doi: 10.1158/1535-7163.MCT-15-0634. PubMed PMID: 28258165; PubMed Central PMCID: PMC5418095.

28. Sviripa V, Zhang W, Conroy MD, Schmidt ES, Liu AX, Truong J, et al. Fluorinated N,N’-diarylureas as AMPK activators. Bioorg Med Chem Lett. 2013;23(6):1600–3. doi: 10.1016/j.bmcl.2013.01.096. PubMed PMID: 23414799; PubMed Central PMCID: PMC3594501.

29. Huang X, Li X, Xie X, Ye F, Chen B, Song C, et al. High expressions of LDHA and AMPK as prognostic biomarkers for breast cancer. Breast. 2016;30:39–46. doi: 10.1016/j.breast.2016.08.014. PubMed PMID: 27598996.

30. Al-Maghrabi J, Al-Sakkaf K, Qureshi IA, Butt NS, Damnhory L, Elshal M, et al. AMPK expression patterns are significantly associated with poor prognosis in breast cancer patients. Ann Diagn Pathol. 2017;29:62–7. doi: 10.1016/j.anndiagpath.2017.05.012. PubMed PMID: 28807345.

31. Hadad SM, Baker L, Quinlan PR, Robertson KE, Bray SE, Thomson G, et al. Histological evaluation of AMPK signalling in primary breast cancer. BMC Cancer. 2009;9:307. doi: 10.1186/1471-2407-9-307. PubMed PMID: 19723334; PubMed Central PMCID: PMC2744705.

32. Khabaz MN, Al-Sakkaf K, Qureshi I, Butt N, Damnhory L, Elshal M, et al. Expression of p-AMPK is associated with hormone receptor phenotypes and lymph node metastasis in breast cancer. International journal of clinical and experimental pathology. 2017;10(6):7044–51.

33. Gwak H, Kim Y, An H, Dhanasekaran DN, Song YS. Metformin induces degradation of cyclin D1 via AMPK/GSK3beta axis in ovarian cancer. Mol Carcinog. 2017;56(2):349–58. doi: 10.1002/mc.22498. PubMed PMID: 27128966.

34. Fingar DC, Richardson CJ, Tee AR, Cheatham L, Tsou C, Blenis J. mTOR controls cell cycle progression through its cell growth effectors S6K1 and 4E-BP1/eukaryotic translation initiation factor 4E. Mol Cell Biol. 2004;24(1):200–16. PubMed PMID: 14673156; PubMed Central PMCID: PMC303352.

